# dadi-cli: Automated and distributed population genetic model inference from allele frequency spectra

**DOI:** 10.1101/2023.06.15.545182

**Authors:** Xin Huang, Travis J. Struck, Sean W. Davey, Ryan N. Gutenkunst

## Abstract

**Summary:** dadi is a popular software package for inferring models of demographic history and natural selection from population genomic data. But using dadi requires Python scripting and manual parallelization of optimization jobs. We developed dadi-cli to simplify dadi usage and also enable straighforward distributed computing.

**Availability and Implementation:** dadi-cli is implemented in Python and released under the Apache License 2.0. The source code is available at https://github.com/xin-huang/dadi-cli. dadi-cli can be installed via PyPI and conda, and is also available through Cacao on Jetstream2 https://cacao.jetstream-cloud.org/.

## Introduction

In population genomics, model-based inference is important for learning about past population history and natural selection (Johri et al. 2022). A common approach is to fit models to data summarized by an allele frequency spectrum (AFS), which is a multi-dimensional array in which each entry counts the number of mutations observed at a given combination of sample frequencies. Multiple approaches exist for inference from AFS (Excoffier et al. 2013; Jouganous et al. 2017; Kamm et al. 2020; Excoffier et al. 2021), and dadi is popular for inferring models of demographic history (Gutenkunst et al. 2009) (population size changes, divergences, and migration) and distributions of mutation fitness effects (DFEs; Kim et al. 2017). Features particular to dadi include computation on graphics processing units (Gutenkunst 2021), uncertainty estimation using the Godambe Information Matrix (Coffman et al. 2016), modeling of inbred (Blischak et al. 2020) and polyploid populations (Blischak et al. 2023), and inference of joint DFEs between populations (Huang et al. 2021). dadi is implemented as a Python library and driven by a script file, which provides flexibility for advanced users but is a barrier for new users. Moreover, parameter optimization within dadi can be computationally expensive and is highly parallelizable, but dadi users must implement their own parallelization framework. To address both these challenges, we implemented dadi-cli, a command-line interface for dadi that supports distributed computation.

## Approach

Through dadi-cli, users can implement the primary dadi workflows, using several subcommands (Fig. 1A). Distributed parallel computing in enabled for the most computationally intense subcommands. For any workflow, a user will typically first use the `GenerateFs` command to calculate the data AFS from an input Variant Call Format file (Danecek et al. 2011).

**Figure 1:**
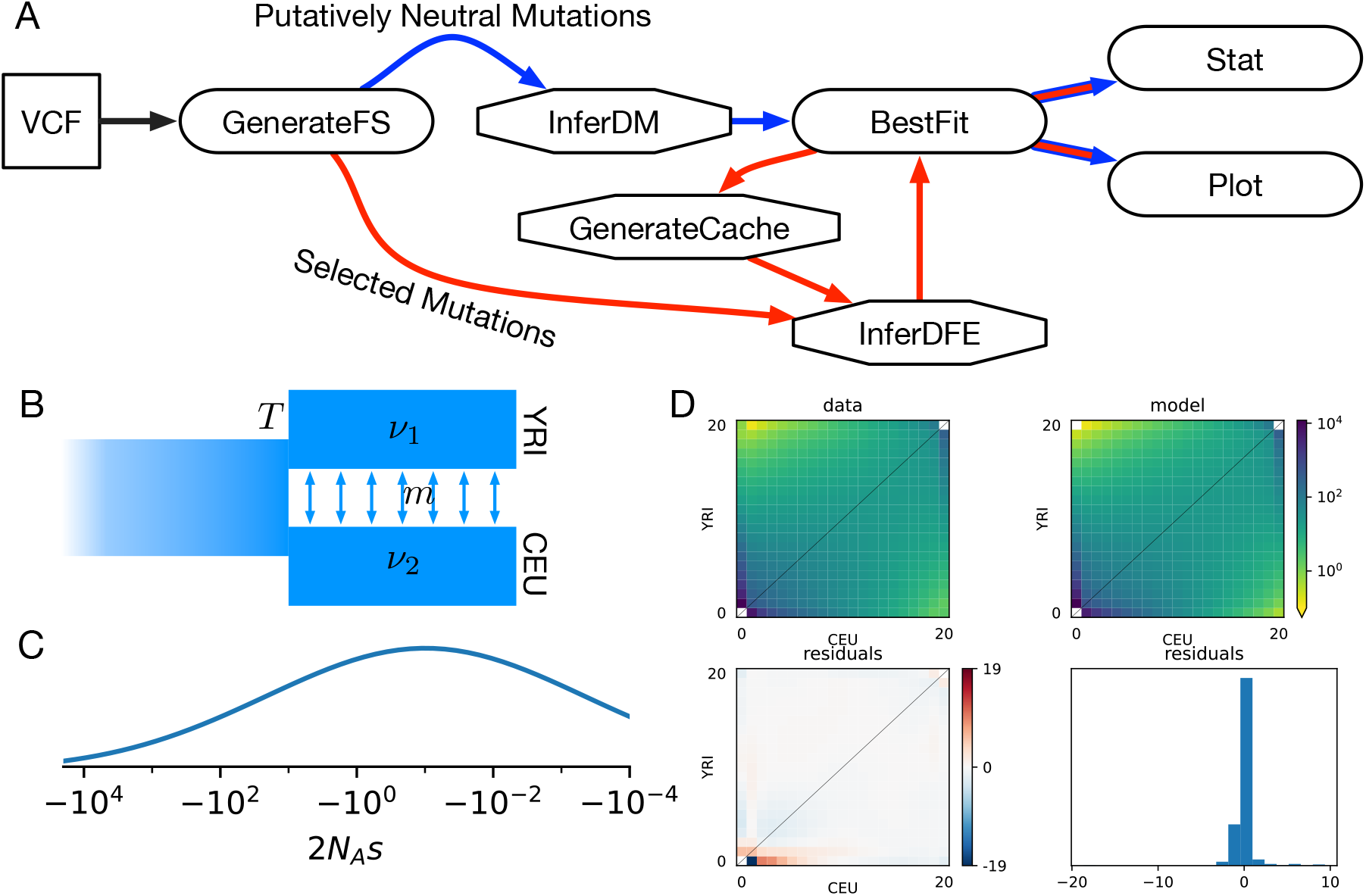
A) Workflows implemented in dadi-cli. Blue arrows indicate a workflow for inferring a demographic model from putatively neutral mutations. Red arrows indicate a workflow for inferring a distribution of fitness effects (DFEs) from selected mutations. Hexagonal nodes indicate subcommands for which parallel processing is implemented. B) Split-migration demographic history model. C) Illustrative lognormal distribution of fitness effects (DFE) of new mutations. D) Model assessment plot. The upper-left panel shows the data allele frequency spectrum (AFS), and the upper-right shows the demographic history plus DFE model AFS. The lower-left panel shows the scaled residuals for each entry in the AFS, and the lower-right shows a histogram of the scaled residuals.

Often the final goal is to infer a model of demographic history, using putatively neutral mutations, such as those in intergenic regions. Arbitrary scenarios of population divergence, size change, and continuous and pulse migration are supported, through models built into dadi or through user-coded custom model functions. A key computational parameter for dadi is the number of grid points used to discretize population allele frequencies. Large sample sizes demand more grid points for accuracy; dadi-cli provides a default calculation sufficient for most demographic inference. Model parameters are optimized to maximize the composite likelihood of the AFS using the `InferDM` subcommand, with the `BestFit` command used to consolidate results among several optimization runs. Once the maximum likelihood parameter set is identified, statistical uncertainties can be estimated using the `Stat` subcommand, and a visual comparison between the model and data can be produced by the `Plot` subcommand.

The DFE of new mutations can be estimated from putatively selected sites, often within coding sequences. To do so, a demographic model is first estimated, typically from synonymous mutations, that corrects for effects of both demographic history and linked selection. Because selected mutations are assumed not to interact, the expected AFS for a given DFE is a weighted sum over spectra for each possible selection coefficient (Keightley & Eyre-Walker 2007). A cache of those spectra is calculated given the demographic model using the `GenerateCache` subcommand. Multiple parametric models for the DFE are supported, like gamma and lognormal distributions, and users can create custom models. The `InferDFE` command is then used to maximize the likelihood of the AFS for selected mutations. The `BestFit`, `Stat`, and `Plot` commands then perform similar functions to the demographic model case. Recently, our group has introduced the notion of a joint DFE between populations (Huang et al. 2021), for which inference is also supported.

Typically, parameter optimization is the most computationally demanding step in dadi inference, implemented in the `InferDM` and `InferDFE` subcommands. Exploring the nonlinear likelihood surface to identify the true maximum likelihood point is challenging. For dadi-cli, we take a multiple-shooting approach, beginning with a short global optimization and then using local optimizations started from many initial parameter sets. In this procedure, convergence is assessed by comparing the likelihoods and parameter values of the best local optimization runs. (See Excoffier et al. (2013) and Noskova et al. (2020) for alternative approaches for exploring parameter space.) It is often unclear how many optimization runs will be necessary to identify the mximimum likelihood parameter values. To aid users, the `--force-convergence` option runs optimizations until convergence criteria are met.

The multiple-shooting optimization approach used by dadi enables straightforward parallelization and distributed computing. When running on a single compute node, dadi-cli uses the built-in Python multiprocessing module to coordinate parallel optimization across local CPU cores. For challenging optimizations, dadi-cli enables coordination across multiple compute nodes through the Work Queue framework within CCTools (Bui et al. 2011). This framework enables communication between a manager process and workers that may be distributed across nodes, coordinated simply by a project name and password file. Typically, users would use a Work Queue factory to automatically spawn workers on each node, while a single dadi-cli process acts as the manager. For users without fixed high performance computing facilities, we supply Terraform scripts that enable users to easily launch Elastic Compute Cloud instances from from Amazon Web Services to run dadi-cli and Work Queue. Lastly, for users unfamiliar with high performance or cloud computing, we have developed a web interface within the Cacao framework to launch jobs on the public Jetstream2 cloud computing system (Hancock et al. 2021).

## Example

As an example, we present and inference of the DFE for nonsynonymous mutations using samples from two human populations, Yoruba in Ibadan, Nigeria (YRI) and Utah residents (CEPH) with Northern and Western European ancestry (CEU), from the 1000 Genomes Project data (The 1000 Genomes Project Consortium 2015). Using BCFtools (Danecek et al. 2021), we first created compressed `.vcf` files containing only the populations of interest and biallelic single-nucleotide mutations in coding sequences. Frequency spectra for synonymous and nonsynonymous mutations can then be extracted into `*.fs` files using the following commands:

~~~
dadi-cli GenerateFs --vcf data/1KG.YRI.CEU.syn.vcf.gz --pop-info data/1KG.YRI.CEU.popfile.txt
   --pop-ids YRI CEU --projections 20 20 --polarized --output results/1KG.YRI.CEU.20.syn.fs
dadi-cli GenerateFs --vcf data/1KG.YRI.CEU.non.vcf.gz --pop-info data/1KG.YRI.CEU.popfile.txt
   --pop-ids YRI CEU --projections 20 20 --polarized --output results/1KG.YRI.CEU.20.non.fs
~~~

(Note that for speed of execution, here we project our frequency spectra down to 20 chromosomes per population, which a complete analysis would avoid.) For initial data quality assessment, a plot of the synonymous AFS can be made using

~~~
dadi-cli Plot --fs results/1KG.YRI.CEU.20.syn.fs --output results/1KG.YRI.CEU.20.syn.pdf
~~~

yielding Fig. S1.

To infer a DFE, we first fit a demographic model to the synonymous mutation data, which accounts for both demographic history and some effects of linked selection. We fit a split with migration model (Fig. 1B) using the following command

~~~
dadi-cli InferDM --fs results/1KG.YRI.CEU.20.syn.fs --model split_mig
  --lbounds 1e-3 1e-3 0 0 0 --ubounds 100 100 1 10 0.5 --force-convergence 10 --cpus 4
  --output results/1KG.YRI.CEU.20.split_mig
~~~

Here the `bounds` arguments define the range of parameter values that will be explored, and the `cpus` argument enables parallel execution of four optimization runs at a time. The four demographic model parameters are the relative sizes of the two populations compared to the ancestral population (*nu*_1_ and *ν*_2_), the divergence time *T*, the migration rate *m*. In addition, by default a parameter is added to account for misidentification of mutation ancestral states (Baudry & Depaulis 2003). The `force-convergence` argument tells dadi-cli to run at least ten optimizations and to continue optimization until the top three parameter sets found are within a narrow range of likelihoods. Note that if the `InferDM` subcommand were run multiple times, the output files would be consecutively numbered, so they could be easily combined using the `BestFit` command:

~~~
dadi-cli BestFit --input-prefix results/1KG.YRI.CEU.20.split_mig.InferDM
~~~

To distribute the optimization runs across nodes using Work Queue, with a project name of `dminf` and a password file `pwfile`, the `InferDM` command could be changed simply by adding the argument `--work-queue dminf pwfile`. Then workers could be spawned by running a factory process on each node, with each worker using a single core

~~~
work_queue_factory -T local -M dminf -P pwfile --cores=1
~~~

The Work Queue framework manages network communication between manager and workers (although crossing firewalls can be a challenge.) Given the optimized parameters, the quality of the fit to the data can be visualized (Fig. S2):

~~~
dadi-cli Plot --fs results/1KG.YRI.CEU.20.syn.fs --model split_mig
  --demo-popt results/1KG.YRI.CEU.20.split_mig.InferDM.bestfits
  --output results/1KG.YRI.CEU.20.syn.vs.split_mig.pdf
~~~

This plot shows that the model poorly fits low-frequency mutations that are private to the CEU sample. This is most likely because this model does not include the exponential growth that is known to have occurred in European populations.

With the demographic model inferred, we now turn to the DFE, assuming that selection coefficients for all mutations are equal in the two populations. We generate a cache of spectra corresponding to different values of the selection coefficient.

~~~
dadi-cli GenerateCache --model split_mig_sel_single_gamma
  --cpus 4 --sample-size 20 20 --grids 280 290 300
  --demo-popt results/1KG.YRI.CEU.20.split_mig.InferDM.bestfits
  --output results/1KG.YRI.CEU.20.split_mig.sel.single.spectra.bpkl
~~~

The caches includes 50 values of the population-scaled selection coefficient between −10^−4^ and −2000. The number of values can be adjusted with `--gamma-pts` and the range can be adjusted with `--gamma-bounds`.

Because model calculation at large selection coefficients in challenging, the `grids` setting is used to enable a finer grid for model integration. Next we maximize the likelihood of the nonsynonymous data by optimize the parameters of a lognormal DFE model (Fig. 1C).

~~~
dadi-cli InferDFE --fs results/1KG.YRI.CEU.20.non.fs --cpus 4
  --cache1d results/1KG.YRI.CEU.20.split_mig.sel.single.spectra.bpkl
  --pdf1d lognormal --lbounds -10 0.01 0 --ubounds 10 10 0.5
  --demo-popt results/1KG.YRI.CEU.20.split_mig.InferDM.bestfits --ratio 2.31
  --output results/1KG.YRI.CEU.20.split_mig.dfe.1D_lognormal --force-convergence 10
~~~

Here the model parameters are the mean and standard deviation of the distribution of log selection coefficients, along with an ancestral state misidentification parameter. An important input is the `ratio` of mutation rates for the creation of nonsynonymous versus synonymous mutations. After optimization, we extract the best fit parameters just as in a demographic history inference

~~~
dadi-cli BestFit --input-prefix results/1KG.YRI.CEU.20.split_mig.dfe.1D_lognormal.InferDFE
~~~

We can assess the quality of our inference both visually and via formal uncertainty analysis. Visually, we plot a comparison of model and the data (Fig. 1D)

~~~
dadi-cli Plot --fs results/1KG.YRI.CEU.20.non.fs --pdf1d lognormal
  --dfe-popt results/1KG.YRI.CEU.20.split_mig.dfe.1D_lognormal.InferDFE.bestfits
  --cache1d results/1KG.YRI.CEU.20.split_mig.sel.single.spectra.bpkl
  --output results/1KG.YRI.CEU.20.non.1D_lognormal.pdf
~~~

We see that the pattern of residuals is very similar to the demographic model fit to the synonymous data (Fig. S2). This is typical in DFE analysis, and the magnitude of the residuals suggests that altering our demographic model may be valuable. We estimate confidence intervals using bootstrap resampling over genomic regions of both our synonymous and nonsynonymous mutations

~~~
dadi-cli GenerateFs --vcf data/1KG.YRI.CEU.syn.vcf.gz --pop-info data/1KG.YRI.CEU.popfile.txt
  --pop-ids YRI CEU --projections 20 20 --bootstrap 100 --chunk-size 1000000
  --output results/boots_syn/1KG.YRI.CEU.20.syn --seed 42
dadi-cli GenerateFs --vcf data/1KG.YRI.CEU.non.vcf.gz --pop-info data/1KG.YRI.CEU.popfile.txt
  --pop-ids YRI CEU --projections 20 20 --bootstrap 100 --chunk-size 1000000
  --output results/boots_non/1KG.YRI.CEU.20.non --seed 42
~~~

Note that we use the random number `seed` argument to ensure that the same genomic regions are chosen in each command. With these bootstraps, we can then run a Godambe Information Matrix analysis, which estimates uncertainties account for linkage between mutations (Coffman et al. 2016)

~~~
dadi-cli StatDFE --fs results/1KG.YRI.CEU.20.non.fs
  --dfe-popt results/1KG.YRI.CEU.20.split_mig.dfe.1D_lognormal.InferDFE.bestfits
  --cache1d results/1KG.YRI.CEU.20.split_mig.sel.single.spectra.bpkl --pdf1d lognormal
  --bootstrapping-nonsynonymous-dir results/boots_non
  --bootstrapping-synonymous-dir results/boots_syn
  --output results/1KG.YRI.CEU.20.split_mig.dfe.1D_lognormal.ci
~~~

We find that the 95% confidence intervals for our estimates of the mean and standard deviations of the logs of the population-scaled selection coefficients are 1.76 − 2.88 and 4.92 − 5.85, respectively. These confidence intervals overlap with the analysis in Huang et al. (2021).

As a command-line tool, dadi-cli is straightforward to use within workflow managers. For example, we provide a SnakeMake (Köster & Rahmann 2012) workflow that fits DFE models to all populations within the 1000 Genomes Project data. As expected, we find similar parameters for all populations (Fig.S3).

In conclusion, dadi-cli is a powerful and convenient tool for population genomic inference. It greatly simplifies the usage of dadi, particularly for complex operations such as uncertainty analysis. Moreover, dadi-cli enables parallel and distributed parameter optimization. As population genomic data become available for more species, dadi-cli will be a useful tool for the research community to explore population history and natural selection.

## Acknowledgements

The authors thank Nirav Merchant and Doug Thain for consultation regarding cloud computing. We also thank the following beta testers: Linh Tran, Connie Sun, Emanuel Fonseca, Andrew Kern, Chris Smith, Will Nash, Niklas Schulmeister, Heng Liang, the dadi-user Google group, and the PopSim Consortium. X.H. thanks the Life Science Compute Cluster (LiSC) of the University of Vienna for support.

## Funding

This work was supported by the National Institutes of General Medical Sciences [R01GM127348 to R.N.G.].

## Author Contributions

R.N.G. and X.H. designed the study. X.H., T.J.S., S.W.D., and R.N.G. implemented dadi-cli. X.H., T.J.S., and R.N.G. wrote the manuscript.

## Competing Interests

The authors declare no conflict of interests.

## Supporting Information

**Figure S1:**
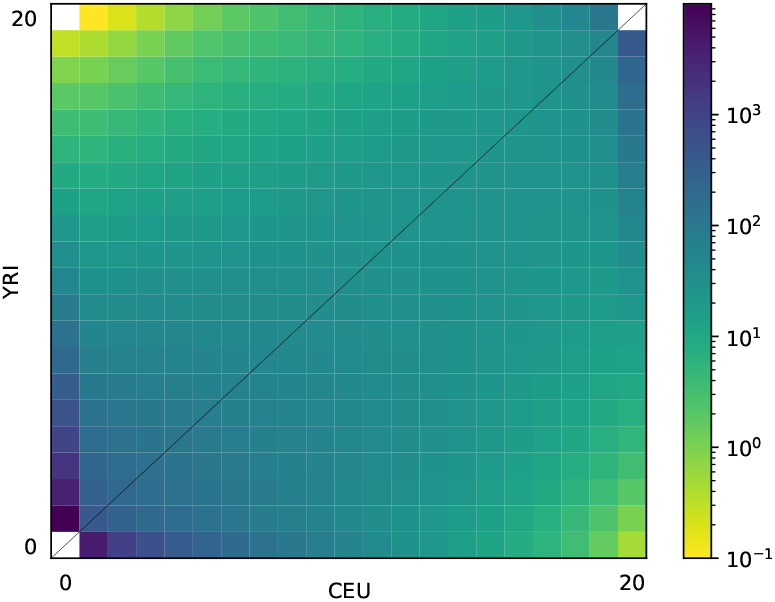
Synonymous AFS extracted from 1000 Genomes Project data.

**Figure S2:**
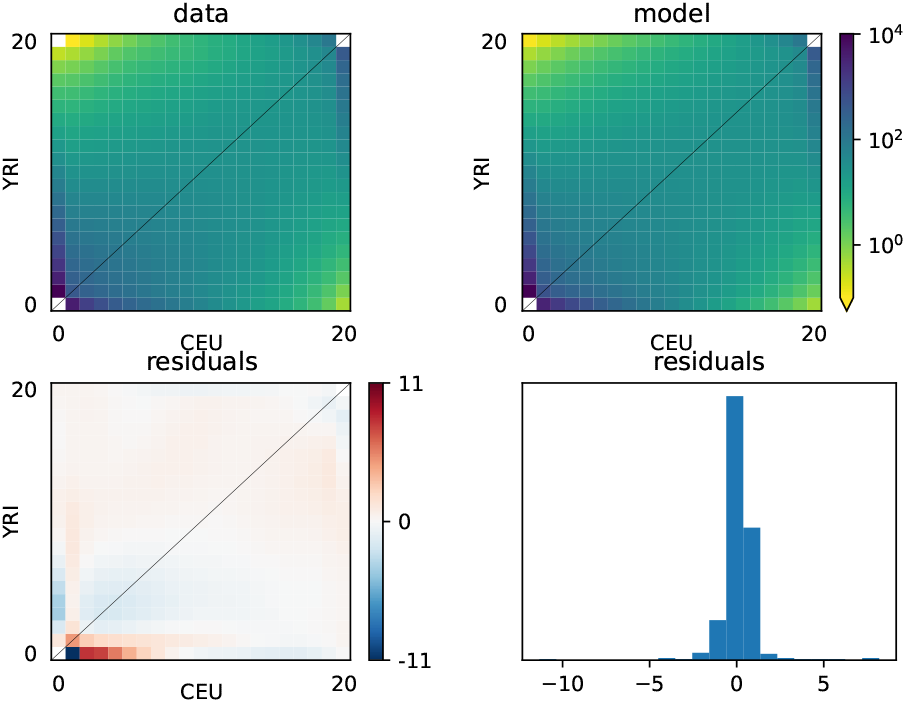
Comparison between synonymous data and split-with-migration demographic model.

**Figure S3:**
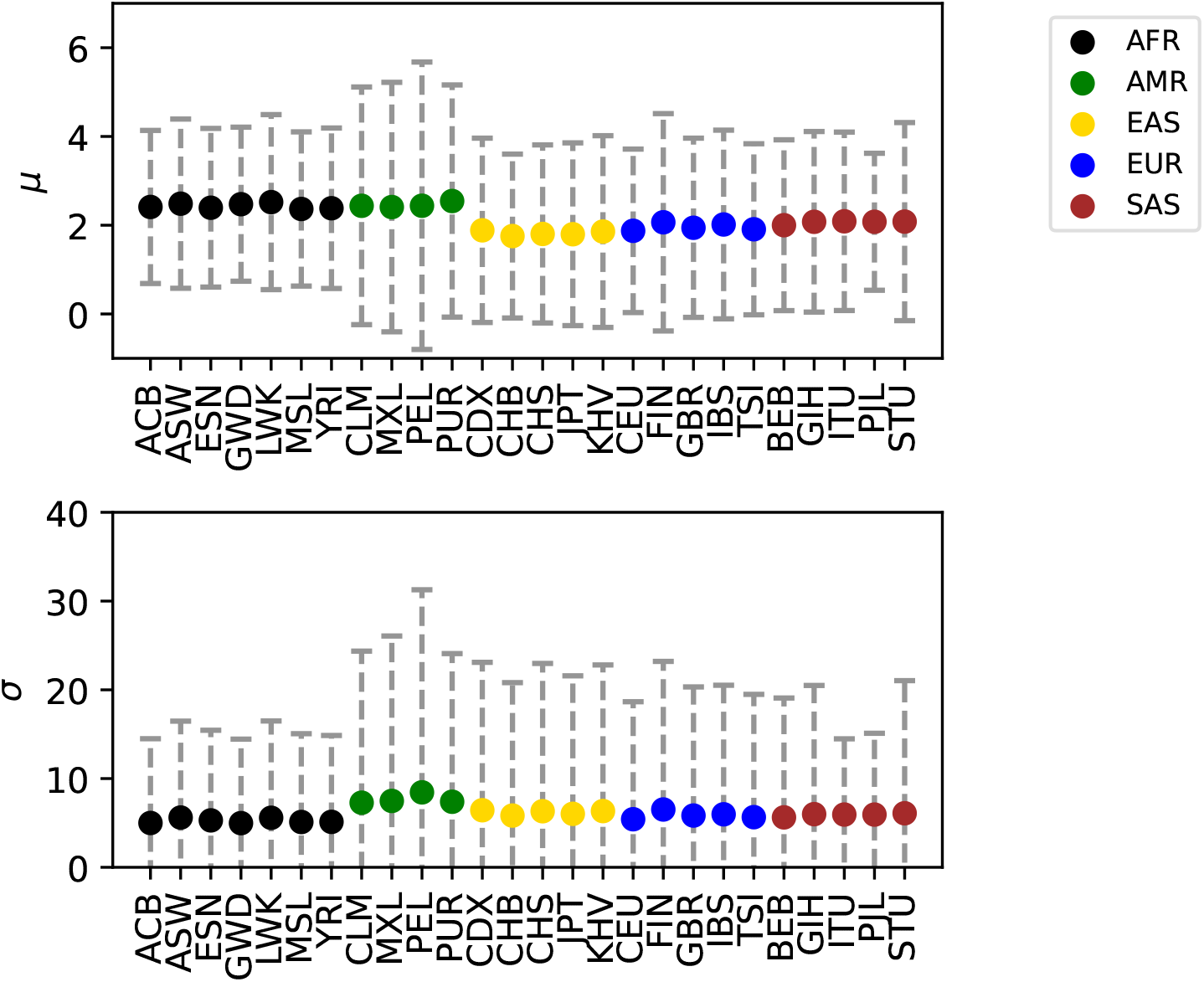
Lognormal DFE model inferences for 1000 Genomes Populations. Top panel shows the mean of the distribution of logs of scaled selection coefficients, while the bottom shows the standard deviation. Whiskers denote 95% confidence intervals. Used abbreviations: AFR, African populations; AMR, American populations; EAS, East Asia populations; EUR, European populations; SAS, South Asia populations; ACB, African Caribbean in Barbados; ASW, African Ancestry in Southwest US; ESN, Esan in Nigeria; GWD, Gambian in Western Division, The Gambia; LWK, Luhya in Webuye, Kenya; MSL, Mende in Sierra Leone; YRI, Yoruba in Ibadan, Nigeria; CLM, Colombian in Medellin, Colombia; MXL, Mexican Ancestry in Los Angeles, California; PEL, Peruvian in Lima, Peru; PUR, Puerto Rican in Puerto Rico; CDX, Chinese Dai in Xishuangbanna, China; CHB, Han Chinese in Beijing, China; CHS, Han Chinese South; JPT, Japanese in Tokyo, Japan; KHV, Kinh in Ho Chi Minh City, Vietnam; CEU, Utah residents (CEPH) with Northern and Western European ancestry; FIN, Finnish in Finland; GBR, British in England and Scotland; IBS, Iberian populations in Spain; TSI, Toscani in Italia; BEB, Bengali in Bangladesh; GIH, Gujarati Indian in Houston, TX; ITU, Indian Telugu in the UK; PJL, Punjabi in Lahore, Pakistan; STU, Sri Lankan Tamil in the UK.

## Notes

https://github.com/xin-huang/dadi-cli

### Competing Interest Statement

The authors have declared no competing interest.

